# Molecular rewiring and compensatory mechanisms sustain DNA recognition in mutant ZTA transcription factor: insights from molecular dynamics simulations

**DOI:** 10.64898/2025.12.17.694912

**Authors:** Boobalan Duraisamy, Debabrata Pramanik

## Abstract

Protein–DNA complexes are stabilized by various interactions forming an interaction network between the protein and DNA molecules. Any change in the system – whether through mutations in the protein or DNA, external factors, or protein conformational transitions -- can alter this interaction network, thereby affecting structural and functional aspects. Employing all-atom classical molecular dynamics, we investigated how the interaction network in the ZTA TF-DNA is rewired when key arginine residues in ZTA are mutated to oppositely charged glutamic acids. Using the MMPBSA technique, we calculated per-residue binding energies for all systems and correlated binding affinity with structural features. Our detailed mechanistic study shows that when key arginine residues are mutated, new interactions are formed either around the mutation site and/or in other ZTA monomer. Through load-sharing, the system attempts to counter-balance the interaction load, leading to reorganization of the interaction network. As the number of mutations increases from single-to-double site, the system is able to partially maintain its structural stability. However, with multi-site mutations, even after reorganization of the interaction network, system cannot sustain its structural stability and therefore becomes destabilized. Despite the structural symmetry of the ZTA TF, we observed asymmetric monomer contributions upon mutation. Overall, our rigorous mechanistic studies provide deeper insights into the mechanism of interaction network reorganization in ZTA-DNA system. These comprehensive insights may be useful for tuning binding affinity and structural adaptability under adverse conditions. Since ZTA is a key factor in the Epstein-Barr virus (EBV), this study will be central to understanding DNA recognition and developing drug therapeutics targeting viral transcription factors in EBV.

## INTRODUCTION

Protein-DNA interactions -- hydrogen bonds, electrostatics, Vander Waals forces etc. -- play a central role in DNA recognition and gene regulatory mechanisms. Interactions providing structural stability to the protein-DNA complexes can be categorized by long-lived stabilizing interactions and short-lived transient interactions. All these interactions together form a stable interaction network in the protein-DNA system. Long-lived interactions are key in maintaining structural stability of the complex. When a mutation is introduced either in the protein transcription factor (TF) or DNA or TF/DNA both, depending on the substitutions site, existing stabilizing interactions get broken, and new transient interactions formed to compensate the key interaction loss. This in turn reorganize the existing interaction network between the ZTA TF and DNA. In a recent study, Hastings *et al.* showed that a mutation in the transcription factor MAX change the way it folds and binds to DNA, making it more selective for certain sequences[1]. Kock *et al.* studied rare natural variants in homeodomain proteins and discovered that single point mutation affects the DNA base recognition[2]. In leukaemia, de la Torre *et al.* showed that mutations in the C/EBPα bZIP domain produce a distinct DNA-binding behaviour, confirming that the interaction network is re-arranged[3]. Mao et al. observed that a single amino acid change in DNA-binding protein can reorganize its hydrogen bonding network and affect how tightly it binds to DNA[4]. Vernon *et al., from* structural and thermodynamic experiments reported that a small residue level substitution can also introduce a change in the shape and hydration at the binding site, forming a new set of DNA interactions[5]. Similar other studies reported the reorganization of the interaction network in various protein-protein[6–8], protein-DNA[9–11] systems and also in the context of allosteric effect in protein-ligand system[12].

Here we studied ZTA transcription factor–DNA system associated with the Epstein-Barr virus (EBV). EBV is one of the cancers causing viruses discovered in 1964. According to statistics, more than 90% of populations are infected by this virus at some stages of their life. The TF ZTA from the viral genome binds to viral DNA in BRLF1 gene and activates the lytic cycle. Any change in transcription factor in the form of mutation can influence the structural and functional features of ZTA TF–DNA complex, thereby affecting the overall ability of the TF in regulating gene expression. Previous experimental studies (Table 1) on ZTA TF showed the effect of mutations and their functional implications. Lee Heston *et al.* performed a complete targeted mutation across all residues (178 to 194) in the DNA binding domain and reported some mutants to be competent whereas some to be defective. Particularly mutating 183, 187, 190 from arginine to glutamic acid are defective in activating Rta, defective in EA-D (the viral DNA polymerase processivity factor), and defective in activating late protein[13]. Ray *et al.* studied the effect of mutations at cysteine 189[14] and asparagine 182[15] with various substituents revealing its diverse impacts on the DNA binding specificity. Further studies revealed that mutations in the DNA binding domain at R179A, Y180E, and A185V not only influence the lytic switch but also disrupt the activation of EA-D [16]. The S186A mutation in Zta reduces DNA binding affinity particularly to methylated sites, impeding lytic cycle transition and downstream the gene expression induction[17–20].

**Table 1:** Previous reported mutation studies highlighting the specific mutation sites and corresponding mutations for the ZTA transcription factor-DNA system. The references have also been provided.

| Sl. no. | Mutation | Authors |
| --- | --- | --- |
| a) | R183K, R187K | Lee Heston *et al.* |
| b) | R179A, R183A, R187A, R190A | Lee Heston *et al.* |
| c) | R179E, R183E, R187E, R190E | Lee Heston *et al.* |
| d) | R179A, Y180E, A185V | Park *et al.* |
| e) | S186A | Zhao *et al.* |
| f) | C189 | Ray *et al.* |
| g) | N182 | Ray *et al.* |

Although numerous experimental and computational studies have investigated the effect of mutations in ZTA TF and its biological consequences, a detailed molecular mechanism underlying the rewiring/reorganization of interactions and the compensatory mechanism of other residues upon mutation remain unexplored. It forms the central focus of this work. In our previous work, we studied the influence of DNA sequences on the ZTA – DNA binding specificity and structural aspects[21]. Following our previous study, we selected ZRE 3 (ZTA responsive element 3) sequence, AAA-TTCGCGA-TGC, among 3 identified ZRE’s, ZRE 1, ZRE 2, ZRE 3 in BRLF1 gene in forming interactions with ZTA TF. Based on previous studies reporting glutamic acid as competent[22, 23], defective[24–31], including transcription factor and DNA binding studies[18,30,35–39], we used glutamic acid as mutant in this study.

Here, we focused on how mutations in the viral transcription factor ZTA influences its interaction with DNA at the molecular level using all-atom molecular dynamics simulations[33, 34]. We computed per-residue binding energy to uncover how individual amino acid substitutions alter the structural stability, binding energetics, and interaction network of the ZTA–DNA complex. To observe the graded effect of mutations, we studied five different systems starting from single to multi-site mutations as follows: system A -- wild type with no mutations; system B -- mutated ZTA at R179E, R183E, R187E, R190E from both monomers; system C – mutated at R183E in B monomer & R190E in A monomer; system D – mutated at R183E in B monomer; system E – mutated at R190E in A monomer (shown in Table 2).

**Table 2:** Various system details studied in this work are provided. Mutation were introduced in monomer A and B of the ZTA protein and details of the substitutions were mentioned.

| System | Protein | DNA sequence<br>(ZRE 3) | Mutation site |
| --- | --- | --- | --- |
| A | ZTA | AAA-TTCGCGA-TGC | No mutation |
| B |  |  | (R179E, R183E, R187E, R190E from monomer A and R179E, R183E, R187E, R190E from monomer B) |
| C |  |  | R190E from monomer A and R183E from monomer B |
| D |  |  | R190E from monomer A |
| E |  |  | R183E from monomer B |

## COMPUTATIONAL METHODS

### System modeling

The ZTA TF-DNA PDB structure was obtained from the Protein data bank (PDB ID: 2C9N). The ZRE 3 sequence of the DNA structure was developed using UCSF software [35] and superimposed on the original structure. Protein structure and missing residues were modelled using SWISS-MODELER[36] (in supplementary information we showed details of the modeled ZTA residues). ZTA protein was mutated using rotamer function in UCSF chimera. We studied five different systems: system A -- refers to the wild type ZTA TF–DNA system; system B --four mutations in monomer A and B both; system C -- double mutations; system D -- single point mutation; system E -- single point mutation (shown in Table 2). In Figure 1, we provided schematic representation showing various mutations and associated five different systems. Figure 2 shows the structural difference between Arginine and Glutamic acid.

**Figure 1:**
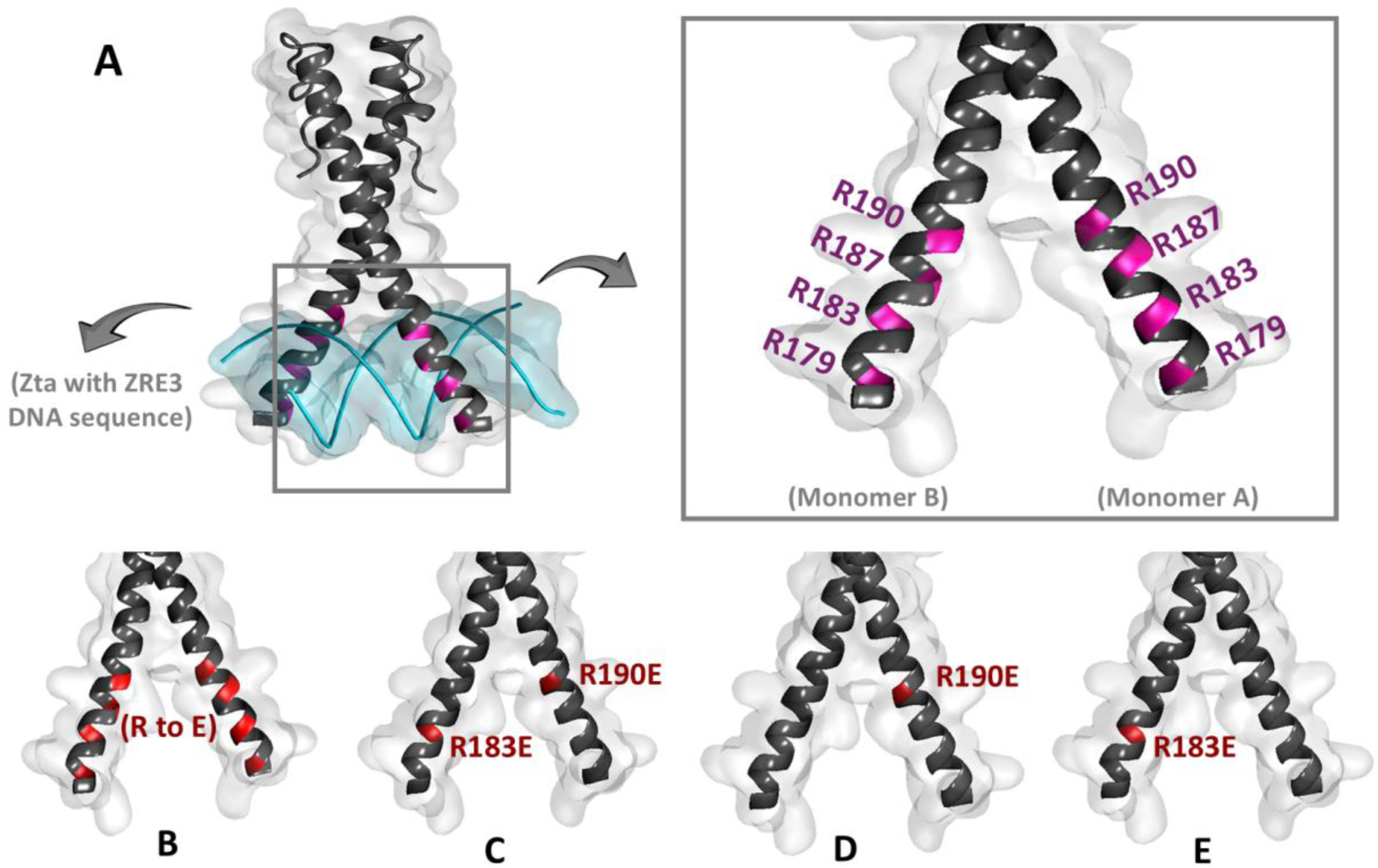
This illustration shows various site-specific mutations studied in this work. (a) System A (Zta protein binds with Viral DNA) wild type system. Inset shows monomer A, monomer B and four arginine residues in each monomer: R179, R183, R187, and R190 respectively. ZRE3 sequence was chosen as DNA sequence, (b) arginine to glutamic acid mutation in all 8 sites in monomer A and B respectively. The substituted sites are shown in red color. (c) ZTA protein is mutated at two sites, R183E from monomer B and R190E from monomer A) (in red). (d) - ZTA mutation at R190E from monomer A. (e) ZTA mutation at R183E from monomer B.

**Figure 2:**
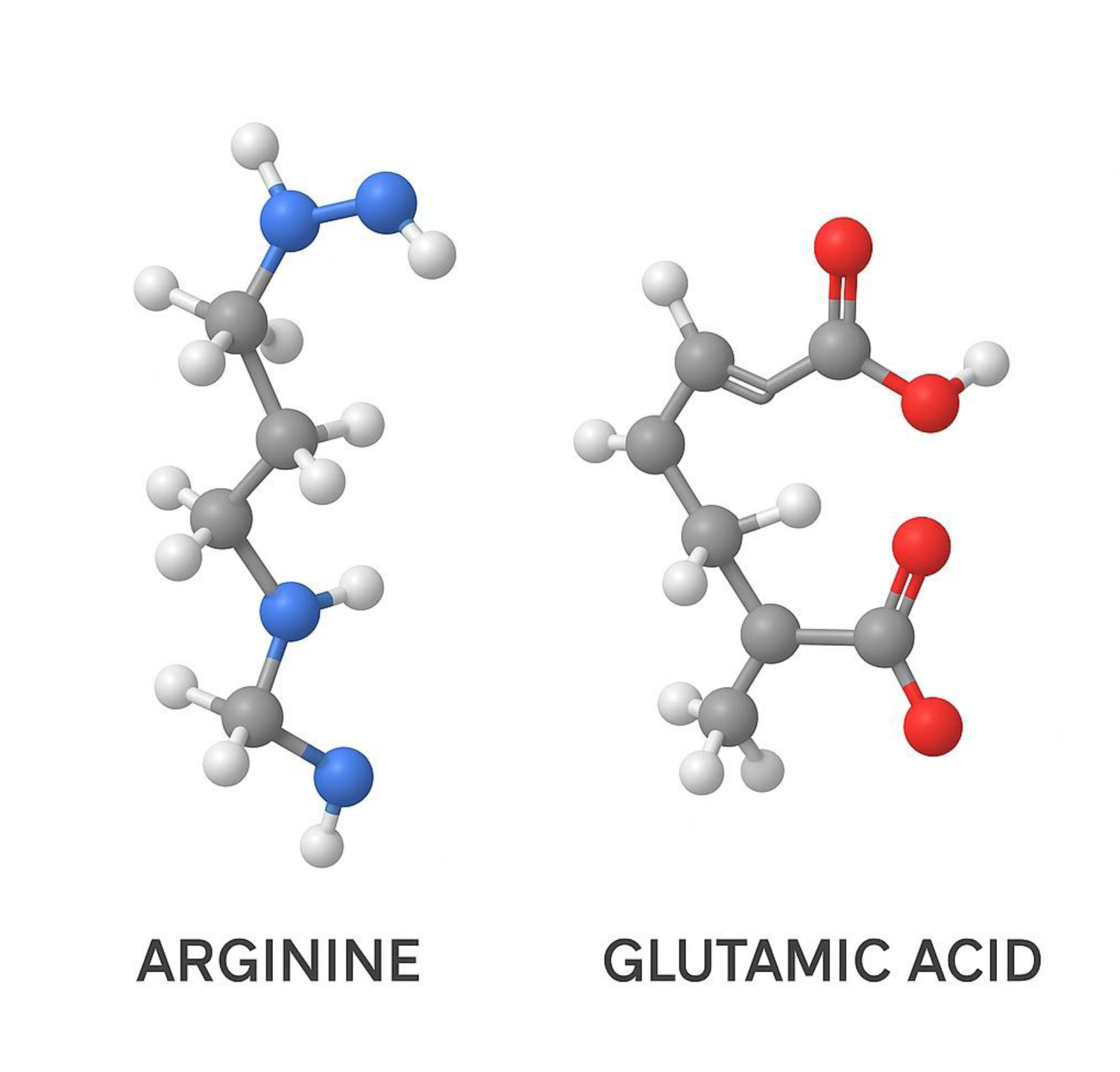
3D optimized structure of arginine and glutamic acid

### Molecular dynamics simulations

To perform classical molecular dynamics (MD) simulations at the all-atom level, we employed GROMACS-2022.4[37] as the basic MD engine. AMBER 03[38] force field parameters were used to model the interaction of protein and DNA systems. TIP3P[39] parameters were used to model the aqueous environment. The systems were charge neutralized by adding Na+ and Cl- counterions. To simulate the bulk properties and remove the finite size effects, PBC (periodic boundary conditions) were applied in all three directions of the system. Once the initial model system was prepared, it was subjected to energy minimization that removes the overlap between solute and solvent molecules. To minimize energy of the system, steepest descent algorithm[40] was employed with a maximum cutoff of 1000 kJ/mol/nm. The neighbour list was updated with every 500 steps (1 ps). To compute various short-range interactions, Verlet[41] cutoff scheme was used with a cutoff distance of 1.4 nm. Followed by energy minimization, the system was equilibrated in NVT and NPT ensemble. For each system, we performed 1 ns of NVT simulation with a time constant of 0.1 ps. The system temperature was maintained at 300 K using v-rescale (velocity rescaling) algorithm. Later, we performed NPT equilibration for 1 ns and the system is maintained with a reference pressure of 1 bar using the Parinello-Rahman Barostat[42] with an isotropic scaling of box vectors. The leap-frog integrator was employed to get the time evolution of the system. Using LINCS [43] algorithm, the hydrogen bonds were restrained which enabled to use an integration time step of 2 fs. The long-range electrostatic interactions were calculated using Particle Mesh Ewald (PME)[44] method with a cubic interpolation order of 4. Once the system reaches equilibrium, we performed production simulation of 500 ns for each system. We analysed these unbiased trajectories to calculate various structural properties and interaction fingerprints. The interaction fingerprints were computed using online available tool ProLIF[45] and VMD[46] software was utilized for visualization of the systems. In Table 3 we provided unbiased simulation details for all five-system studied.

**Table 3:** Various systems studied, box sizes, total number of atoms in the system, number of water molecules, number of counterions and total simulation time are provided for each system for the unbiased MD simulations.

| System | Box sizes (nm<sup>3</sup>) | Total no. of atoms | No. of water molecules | No. of ions | Simulation time (ns) |
| --- | --- | --- | --- | --- | --- |
| A | 12 x 8 x 12 | 113637 | 36887 | 144 | 500 |
| B | 12 x 8 x 12 | 113548 | 36879 | 160 | 500 |
| C | 12 x 8 x 12 | 113884 | 37020 | 140 | 500 |
| D | 12 x 8 x 12 | 113903 | 37024 | 144 | 500 |
| E | 12 x 8 x 12 | 113912 | 37027 | 144 | 500 |

### Binding energy decomposition

To calculate the per-residue free energy for the ZTA protein to identify the key residues contributing to the interaction between ZTA and DNA, we decomposed the overall binding energy. The decomposition analysis was performed using MM/PBSA tool[47]. Binding energy of the complex (ΔGbinding) is calculated using the following equation (eq: 1):

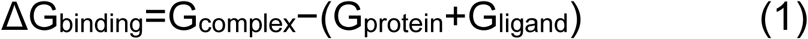

Where, Gcomplex, Gprotein, Gligand – are energies of the complex, only protein, only ligand, respectively.

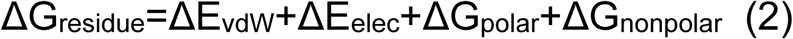

Per-residue binding free energy contributions (ΔGresidue) were calculated using eq(2). Here, ΔEvdW, ΔEelec, ΔGpolar, ΔGnonpolar -- corresponds to VdW, electrostatic, polar and non-polar contributions, respectively. To compute binding energy for each of the system, we selected only last 100 ns data from total 500 ns unbiased simulation trajectories. From 100 ns data, all frames were taken for calculations. The Poison Boltzmann implicit solvent model was employed to calculate the solvation free energy with an ionic strength of 0.15 M. From binding energy decomposition analysis, we computed individual residue contribution to the total binding free energy that allowed us to identify residues with favourable or unfavourable energetic contributions.

### Structural and interaction analysis

To investigate structural stability of the ZTA TF – DNA complexes, for each system we analysed MD simulation trajectories of full 500 ns simulation. To characterize structural stability, we computed RMSD (Root mean square deviation), RMSF (Root mean square fluctuation), and centre of mass (COM) distance using GROMACS[37] in-built packages. In addition, principal component analysis (PCA) was performed to capture the dominant conformational motions of the complexes. The first two principal components (PC1 and PC2) were extracted to describe the major collective movements and to compare the conformational space sampled by the wild-type and mutant systems. Persistent protein-DNA interactions (hydrogen bonds, Van der Waals, pi-pi interaction, cationic and anionic interaction, electrostatic interaction and so on) were computed using the ProLIF[45] tool. In supplementary information we provided details of RMSD, RMSF and various interaction criterion for all kind of interactions (hydrogen bonds, Van der Waals, pi-pi interaction, cationic and anionic interaction, electrostatic interaction) studied. Also, Table S1(a-e) shows details of interaction fingerprints including all kinds of interactions present for all five systems.

### Time variation of hydrogen bonds and contact maps analyses

For each system, two sets of analyses were performed to characterize the ZTA–DNA interactions. All possible non-covalent contacts were first calculated from the last 100 ns unbiased MD trajectories using ProLIF tool after removing translational and rotational motions for proper alignment. For these interactions, two types of plots were generated: (i) a time-series barcode plot showing the temporal evolution of contacts throughout the trajectory, and (ii) a heat map representing the quantified contact frequencies across all residues. The same approach was applied specifically for hydrogen bonds, providing both time-resolved and occupancy-based visualizations. Together, these analyses highlight the dynamics and stability of overall contacts and hydrogen bonds across all five systems (A–E). The interaction cutoff parameters used in ProLIF, and time series bar code plots were added in the Supplementary information (Figure S1 (a-e)).

## Results and discussion

### Mutation sites selection

Following previous mutational studies on ZTA TF-DNA complex (shown in Table 1) and a recent study by Duraisamy et al.,[21], we selected arginine residues to mutate -- R179, R183, R187 and R190 -- from monomer A and monomer B, respectively. From binding energy decomposition analysis, we computed per residue binding energy contributing to the total binding energy for the wild type ZTA TF-DNA (system A) (shown in Figure 3). Figure 3(a) heat map demonstrates binding energy contributions for arginine (ARG) and lysine (LYS) residues across monomer A and monomer B. A deeper blue shade represents minimum binding energy of approximately – 18 kcal/mol. Residues R190 from monomer A and R183 from monomer B show lowest binding energies, approximately – 12 kcal/mol and – 18 kcal/mol. It demonstrates that R190 from monomer A and R183 from monomer B play crucial role in the interaction network and stabilizing ZTA TF-DNA complex. Figure 3 (b) represents schematics of ZTA TF – DNA complex with arginine and lysine residues. A comparative analysis shows total binding energy contribution to be higher for monomer B, approximately – 80 kcal/mol, with respect to monomer A, approximately – 72 to -74 kcal/mol (Figure 3 (c)). Based on these results, we selected R190 in monomer A and R183 in monomer B as mutation targets for further investigations -- systems C, D, E.

**Figure 3:**
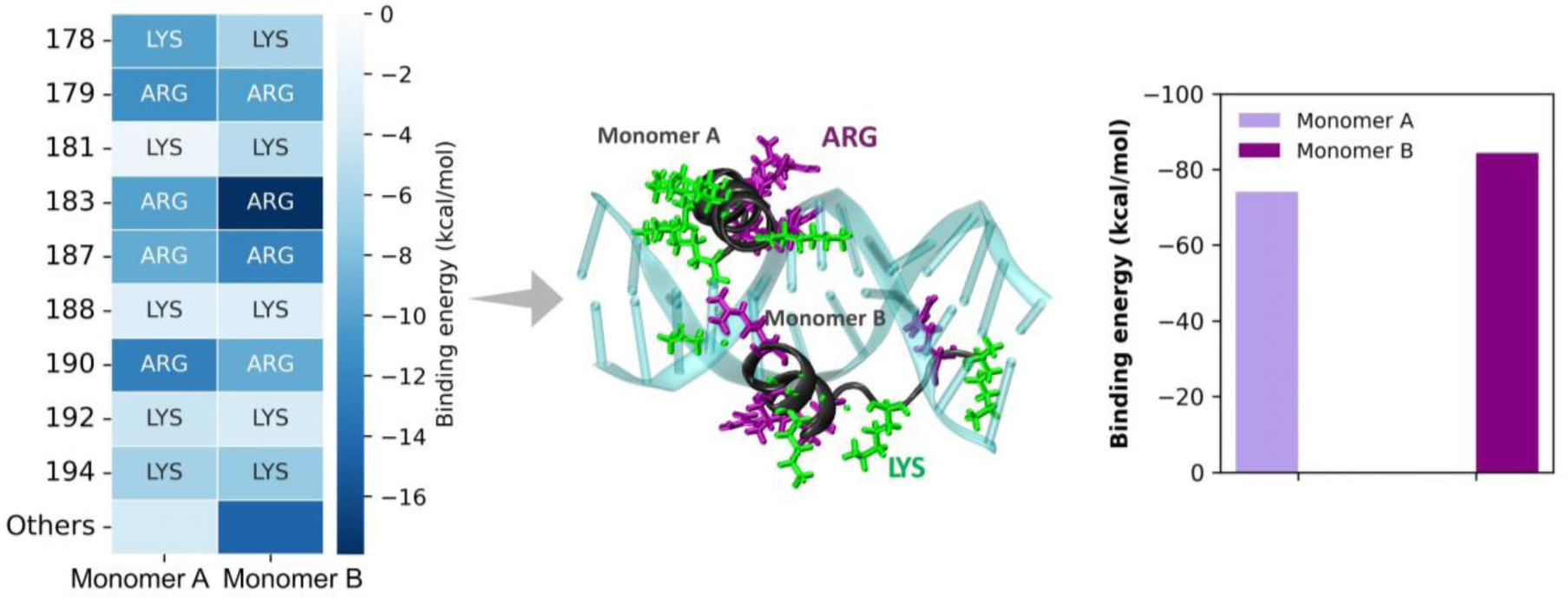
The illustration shows (a) decomposition analysis of binding energy of system A, wild type ZTA – DNA system with the contribution of each monomer. (b) schematic represnetation of the ZTA TF (protein backbone in black) – DNA (in transparent green) complex with arginine residues in violet colour and lysine residues in green colour. Monomer A and B are also mentioned. (c) total binding energy pre monomer A and B respectively..

### Effects of mutations on the structure of ZTA-DNA complex

To explore the effect of arginine substitutions by glutamic acids on structural stability of the ZTA-DNA complexes, we computed RMSD (root mean square deviation), RMSF (root mean square fluctuation), and COM (center of mass) distance for each system. In wild type ZTA TF-DNA system (system A), protein and DNA RMSD (Figure 4(a-b)) remain constant throughout the simulation with minor fluctuations at the starting of the simulation. RMSF analysis shows minimal fluctuations in both the DNA chains and ZTA protein (Figure 4(c) and Figure 5(a)). The COM distance remains constant (approximately 2.5 - 3.0 nm) throughout the simulation (Figure 6). These analyses show overall stability of the wild type ZTA-DNA complex over the simulation time. In system B, we mutated four-sites: R179E, R183E, R187E, R190E, respectively, both in monomer A and B. The protein and DNA RMSD remain almost constant with minor fluctuation over the simulation time (Figure 4(a-b)). The RMSF revealed larger fluctuations in specific DNA nucleic bases between positions 5 and 10, in comparison to other un-mutated regions (Figure 4(c)). The protein RMSF (Figure 5 (b)) shows minor changes around the mutation sites reflecting the effect of mutations on structural rigidity. The COM analysis shows higher fluctuations indicating lesser stability in comparison to other systems (Figure 6). Thus, we observe that multi-site mutations, specifically substitutions of arginine residues, which form key interactions with DNA at the DNA binding domain, cannot preserve structural integrity and stability.

**Figure 4:**
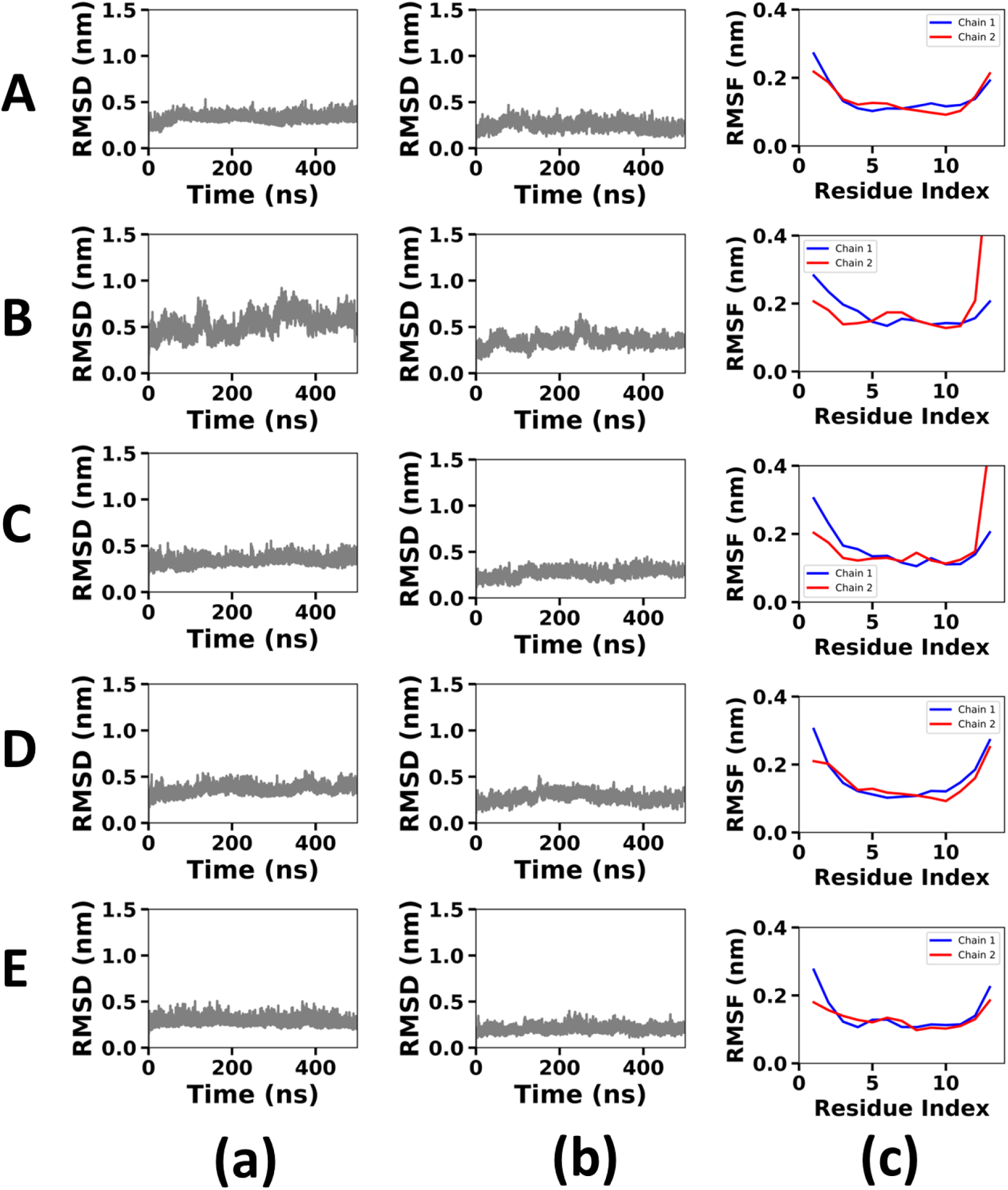
The plot shows time evolution of RMSD (root mean square deviation) over 500 ns of time for (a) protein and (b) DNA systems. Figure (c) shows RMSF (Root mean square fluctuation) of DNA chains in blue and red respectively for chain 1 and chain 2.

**Figure 5:**
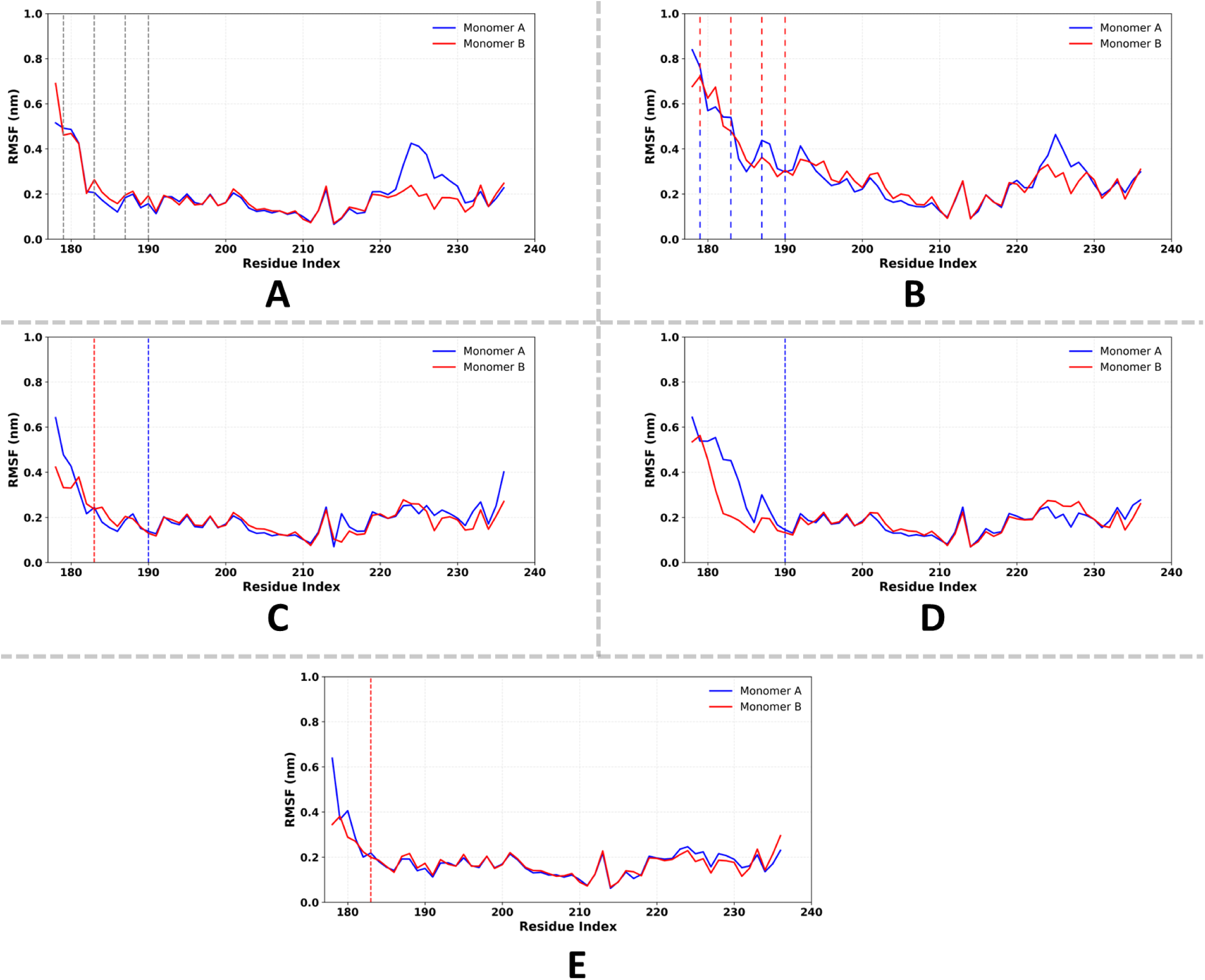
The plot shows RMSF (Root mean square fluctuation) of protein monomers, monomer A (blue) and monomer B (red) respectively. (A) RMSF for wild type ZTA protein. The grey dotted lines show the four arginine residues position in both the monomers where mutation were introduced in later systems. (B) ZTA system with four mutations in each monomer. The blue dotted line shows respective positions of mutation in Monomer A and red dotted line in Monomer B. (C) ZTA system with one mutation in each Monomer. (D) ZTA system with single point mutation in Monomer A. (E) ZTA system with single point mutation in Monomer B.

**Figure 6:**
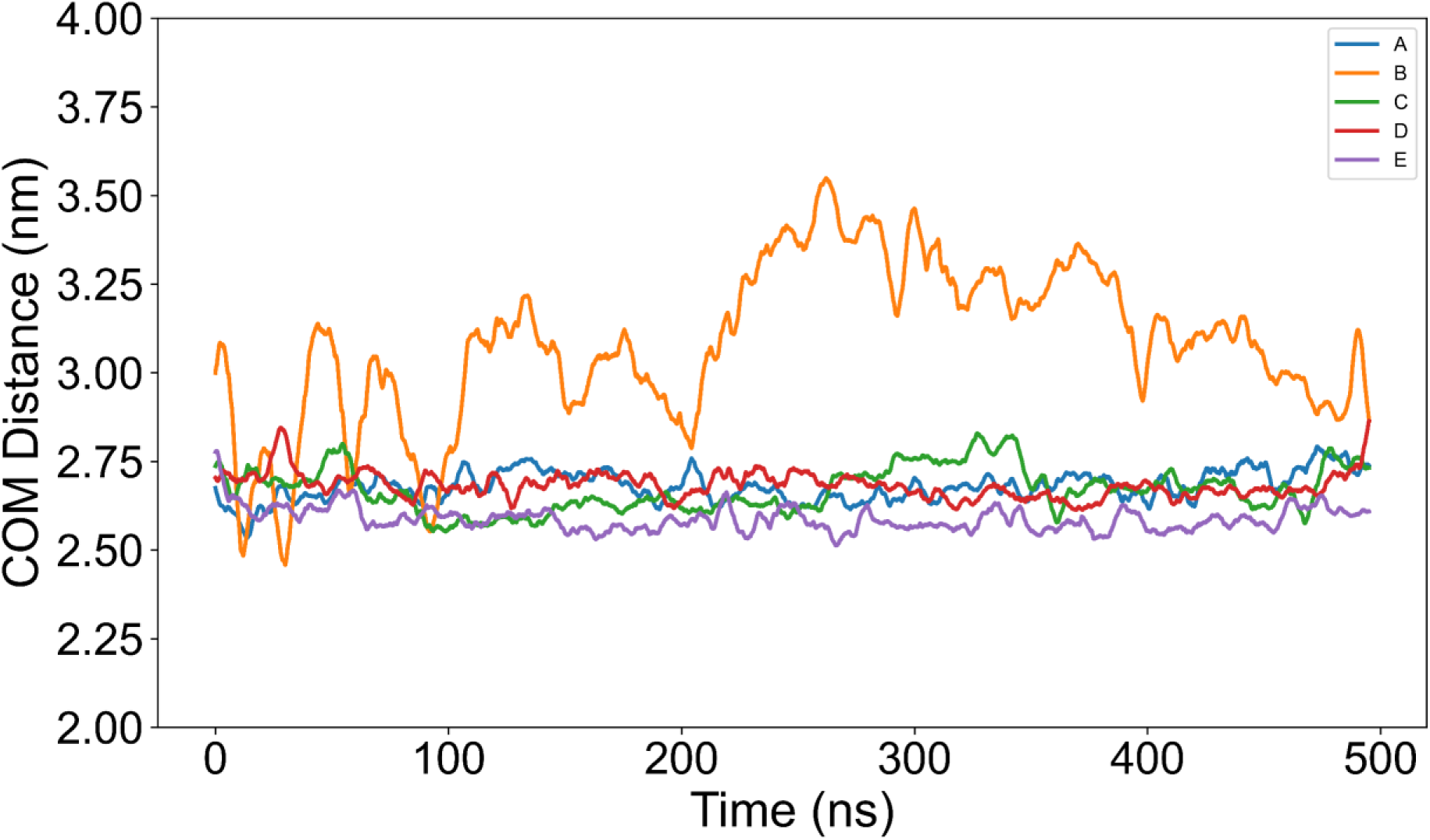
Time evolution of distances between the center of masses of ZTA protein and DNA for all five systems. System A COM distance is represented in blue color, system B in orange, C in green, D in red and E in violet color respectively.

In system C, we mutated two-sites: R183E in monomer B & R190E in monomer A. Protein and DNA RMSD remain almost stable throughout the simulation (Figure 4(a-b)). The DNA RMSF shows fluctuations in residues 5-10 (Figure 4(c)). However, protein RMSF (Figure 5(C)) and COM distance (Figure 6) mostly remain consistent throughout the simulation. Therefore, due to double-site mutations, ZTA-DNA complex can maintain its overall structural stability with minor structural flexibility. In system D and E, a single point mutation was performed at R190E in monomer A and at R183E in monomer B, respectively. Protein and DNA RMSD, protein and DNA RMSF, and COM distance plots show minor fluctuations in both these systems. The structures remain mostly stable throughout the simulation, suggesting structural stability in both systems. Moreover, we observe that a single-point mutation in monomer A at site R190E increases the fluctuation around this residue along the same monomer, whereas monomer B remains relatively stable. Similarly, while mutation is at R183E in monomer B (system E), residue-level fluctuations are higher around the N-terminal region of monomer B, whereas monomer A remains unaffected. This shows that even a single-point mutation to key residue can disturb the symmetry and coordination between monomers in ZTA.

In Figure 5(A-E), we showed residue level analysis of ZTA RMSF and mutation sites by vertical dashed lines/vertical solid lines. In system A which represents wild type, the residues R179, R183, R187, R190 highlighted by vertical dashed lines exhibiting moderate flexibility and symmetrical fluctuations in both monomers, indicating an effective interaction and coordination between monomers. In contrast, system B, with multiple mutations, an increased flexibility is observed between monomers. System C includes double point mutations, monomer B showing slightly reduced flexibility in the mutated region compared to monomer A. This suggests that the mutation in monomer A results in enhanced local fluctuations.

Thus, structural analyses show that with single and double-point mutations, ZTA-DNA complex maintains its partial structural stability with minor fluctuations whereas with multi-site mutations, structure gradually losses its rigidity and become unstable.

### Variation in structural compactness due to mutations

To examine the effect of mutating key arginine residues by glutamic acid on the conformational landscape of ZTA-DNA complex, we performed principal component analysis (PCA) on unbiased molecular dynamics trajectories. We calculated movement of the conformational landscapes along the first two principal components -- PC1 and PC2. The 2D projections of PC1 and PC2 are shown in Figure 7. Each point in the plot refers to a conformational snapshot, and the density of point shows frequency of sampling of those conformations. System A exhibits a narrow and compact distribution centred around the origin indicating a minimal conformational fluctuation (Figure 7(a)). In contrast, system B shows a scattered distribution of points with two major conformational states (Figure 7(b)) – implying a weaker interaction between ZTA and DNA. System C presents a moderately elongated distribution along a diagonal axis, reflecting intermediate flexibility. System D displays a symmetric and slightly expanded distribution compared to A, suggesting a moderate flexibility. System E reveals relatively compact distribution, implying constrained motion with a preferred directionality along its principal components. Thus, PCA analysis reveals that moving from wild type to single-to-double point mutations, the ZTA-DNA complex partially maintain its structural compactness whereas with multi-site mutations to key arginine sites disrupt the cooperative dynamics between monomers leading to a loss in structural stability.

**Figure 7:**
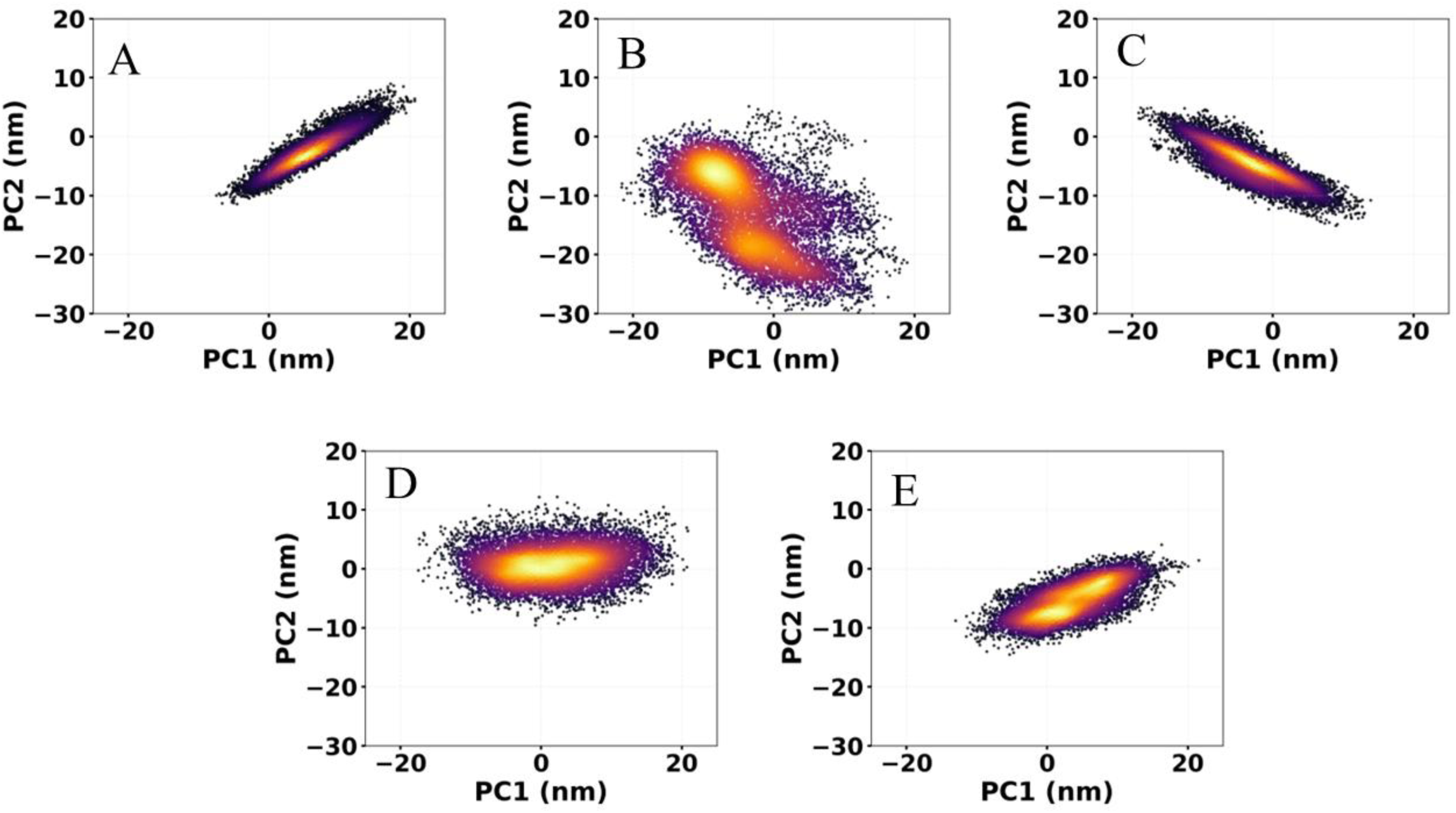
Principal component analysis (PCA) for all five systems. The plot shows PC1 vs PC2 plots for all systems at the bound state of ZTA protein and DNA.

### Contact map analysis: a comparative interaction network rewiring across systems

To analyse the effect of mutating key arginine residues by glutamic acid on ZTA-DNA interaction network, we calculated contact map for all systems (Figure 8). The contact map analysis count the contacts between protein monomers and DNA strands. For DNA, the first 13 nucleic bases (1 to 13) correspond to strand I and 14 to 26 nucleic bases, complementary sequences, form strand II. Each cell color intensity represents the percentage of total contacts considering all kinds of non-covalent interactions. In system A, monomer B shows higher occupancy with strand I, in comparison to monomer A. The arginine residues, B_R179, B_R183, B_R187 and B_R190 form a clear contact band in this region highlighting the key DNA identifying residues. Additionally, B_S186, B_K194, B_V184 and B_N182 residues also show higher percentage of occupancy. On the other way, B_K178, B_R179, B_Y180, B_K192 residues are observed to form transient interactions with strand II. Monomer A behaves in a complementary manner forming majority of the contacts with strand II, the opposite side of the duplex. The arginine residues A_R179, A_R183, A_R187, A_R190 form a parallel strip of most consistent contacts providing stability to that strand. Other residues A_K178, A_Y180, A_N182, A_K194 formed interactions with strand II. A_K188, A_C189, A_K192 forms transient interactions with strand I. Thus, wild-type ZTA–DNA complex shows a directional, cooperative binding mode where monomer B recognizes the core motif on strand I, while monomer A identifies complementary strand II. Roughly 60–65 % of all persistent contacts are formed by arginine residues, 25–30 % from lysines, and less than 10 % from other side chains. About 70 % of all contacts occur on each monomer’s preferred strand (B → I and A → II). This pattern defines the reference interaction layout of the wild-type complex, where the two monomers act together like a clamp—one reading the DNA sequence and the other supporting the duplex for overall stability.

**Figure 8:**
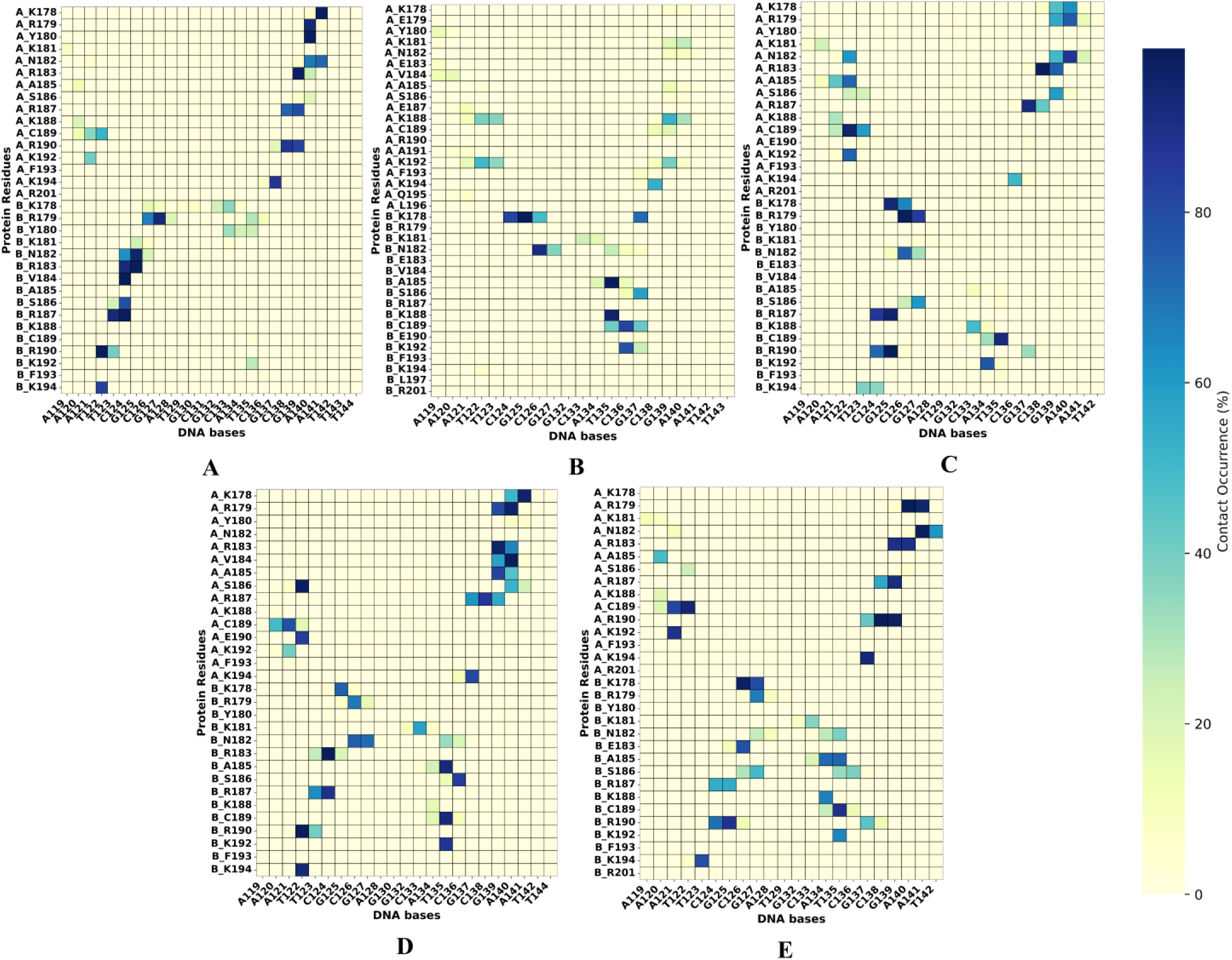
Contact frequency maps for five different protein–DNA systems (A–E), including both wild-type and mutant complexes. The color intensity represents the percentage of total contacts (encompassing hydrogen bonds, hydrophobic, π–π, and van der Waals interactions) formed during molecular dynamics simulations. Compared to the wild-type (A), the mutant systems (B–E) exhibit distinct contact distribution patterns, indicating that mutations alter both the strength and geometry of the protein–DNA interface, potentially modulating overall binding stability.

For system B, the contact map shows a distinct loss of persistent interactions as observed in system A. Most of the contacts formed are transient and scattered. Although both the monomers have been mutated symmetrically, monomer B forms most detectable contacts, specifically B_K178. It interact over most of the simulation time. Nearby residues—B_N182, B_A185, B_K188, B_C189, B_K192 form interactions in both -- strand I and II, whereas monomer A contributes little to the interaction. Only short-lived contacts appear around A_K188, A_K192, and A_K194, mostly on strand II. Overall, system B shows an uneven interaction profile dominated by monomer B. The two-strand connectivity observed in system A collapses to a single-sided pattern centered on B_K178, which provides the strongest and most persistent contact with the DNA core region.

In wild type, we observed monomer A to form interactions predominantly with strand II, whereas in system C, monomer A forms interactions with both strands. Similar interaction pattern is observed for monomer B as well. Hence, upon double-point mutations, the interaction network gets re-distributed and cross-strand contacts are observed which is different from the wild type network pattern. Surrounding residues -- A_C189 and B_C189—nearby the mutant site, respectively form major contacts with strand I and II. In addition, A_K192 forms a new localized transient interaction. The same region on strand I is now occupied by overlapping interactions from B_K178 and B_R179, indicating both residues form contact with the adjacent bases. Apart from the above mentioned residues, lysine residues – for example A_K178–A_K178 and several other lysines from monomer B -- form major interactions as shown in the contact map with broader, spread-out contact blocks. Also, residue -- B_V184 – which previosly form contact with higher strength in wild type, now it is not forming any inetarction. Altogether, these mechanistic details refer to the rewired/reorganized interface: contacts are re-distributed between and across strands, with multiple residues side by mutation sites on the same monomer and/or opposite monomer, stepping in to maintain bonding in case of original contacts are reduced.

In system D, mutation site: A_R190E forms only a single contact with T122. Contrary, R187 interacts with several neighboring bases. A_S186 and A_C189 residues are observed to interact with strand I & II. Many contacts that were previously associated with monomer A, appeared to shift to monomer B; several residues on monomer B (around positions: 185, 186 and 189) and K192, now show markedly stronger interactions with strand II, and additional B-side lysines (including the residues at positions 194 and 178) play key role in forming prominent contacts. Particularly, S186 displays a very strong, persistent contact with nucleic base T122, as a leading interaction compensator. Altogether, the contact map indicates the substitution of A_R190E is compensated by neighboring residues, with an increased engagement from monomer B towards strand II, producing a clear rewiring of contacts between and across the two strands.

Similarly, in system E, the substituted residue B_R183E forms only a single contact with C126, and the earlier strong interaction in wild type at this position is now lost. Several neighboring residues surrounding the mutation site in monomer B -- R179, N182, S186, R187, R190 --now show very weak or almost negligible strength, indicating that most of the pre-existing strong contacts on strand I have faded away. The residue B_K178 remains one of the few strongly interacting positions in monomer B, showing a continuous and intense contact along strand I. Simultaneously, multiple strong interactions have now shifted towards monomer A due to mutaton at B_R183. Distinct and persistent contacts are now seen from A_K192 and A_C189, both showing well-defined patches on strand II. In addition, the arginine residues of monomer A (A_R179, A_R183, A_R187, and especially A_R190) show extended and stronger contacts in comparison to other systems, producing a prominent band across several adjacent bases on strand II. Overall, the map indicates that the loss of interactions around B_R183E leads to a shift of strong DNA contacts from monomer B to monomer A. Especially with K192 and C189 from monomer A, together with multiple arginines, form the principal binding interface in this mutant system, notably monomer A now extend strong interaction towards strand I as well.

The overall structural and interaction analyses reveal how mutations at key arginine residues reshape the stability and recognition dynamics of the ZTA–DNA complex. The wild-type system (A) shows a stable, symmetrical binding pattern where both monomers interact cooperatively with opposite DNA strands. Arginine residues dominate the interaction network, particularly along the guanine-rich core motif, producing strong hydrogen and van der Waals contacts that maintain compact conformations. In contrast, the multiple-mutant system (B) exhibits a significant loss of contacts and structural coordination. Hydrogen bonds and Van der Waals interactions decrease sharply, RMSF fluctuations increase, and PCA shows multiple conformational states — all reflecting weakened DNA binding due to the removal of essential arginine residues. The double mutant (C) displays partial recovery of interaction balance. Contacts are redistributed across both strands, with residues such as A_C189 and A_K192 forming new or extended interactions and lysines contributing to backbone stabilization. This adaptive redistribution aligns with its moderate RMSD and PCA results, showing local flexibility but global stability. The single-point mutants (D and E) demonstrate a clear compensatory behavior. When one monomer is mutated, the other strengthens its DNA contacts — monomer B dominates in system D (mutation in A_R190), while monomer A compensates in system E (mutation in B_R183), even surpassing the wild-type in overall contact count. In both cases, lysine residues and neighboring arginines expand their roles, maintaining hydrogen-bond and ionic interactions that sustain duplex stability. Overall, these findings highlight a robust counter-balancing mechanism within ZTA: when arginine-based contacts weaken on one side, nearby residues and the opposite monomer reorganize their interactions to preserve overall DNA binding and structural integrity. The combination of stable RMSD profiles, localized RMSF fluctuations, and rewired but persistent contact maps indicate that while multiple substitutions disrupt the system severely, single or paired mutations are effectively buffered through adaptive redistribution of interactions. This dynamic compensation between the two monomers underscores the cooperative and resilient nature of the ZTA–DNA binding interface.

### Quantitative analyses of overall interaction change due to mutations

How the total number of interactions, interaction types, key ZTA residues, and crucial nucleic bases are re-distributed upon mutation is important in analysing the detailed mechanistics of ZTA-DNA binding. While computing the interaction, we considered only interactions staying for more than 70% of the total simulation time. Figure 9(a) shows averaged total interaction counts including all types of intercations between ZTA TF and DNA -- 25 (system A), 10 (system B), 14 (system C), 21 (system D), and 32 (system E). We observed total interaction counts to decrease with the number of mutations: no mutation (system A) > single site mutation (system D) > double sites mutation (system C) > multi-site mutations (system B). The only exception was with system E, a single site mutation having higher interaction counts in comparison to zero mutation. Although both, system D and E mutated at single point, however, the total interaction counts differ pointing to the monomer- and site-specific asymmetric behaviour in ZTA-DNA system. A deeper analysis showed that while monomer A is mutated (system D), the interaction counts decreases in monomer A nearly to zero, whereas counts increases largely in monomer B. Contrary, while monomer B is mutated(system E), interaction counts in monomer B remains mostly similar in comparison to wild type, whereas in monomer A, counts increases to a higher value. Hence, ZTA TF whch is structurally symmetric, mutation at key site in one monomer transfers the interaction load to other monomer to counter-balance the load and thus maintain structural stability of the ZTA-DNA complex.

**Figure 9:**
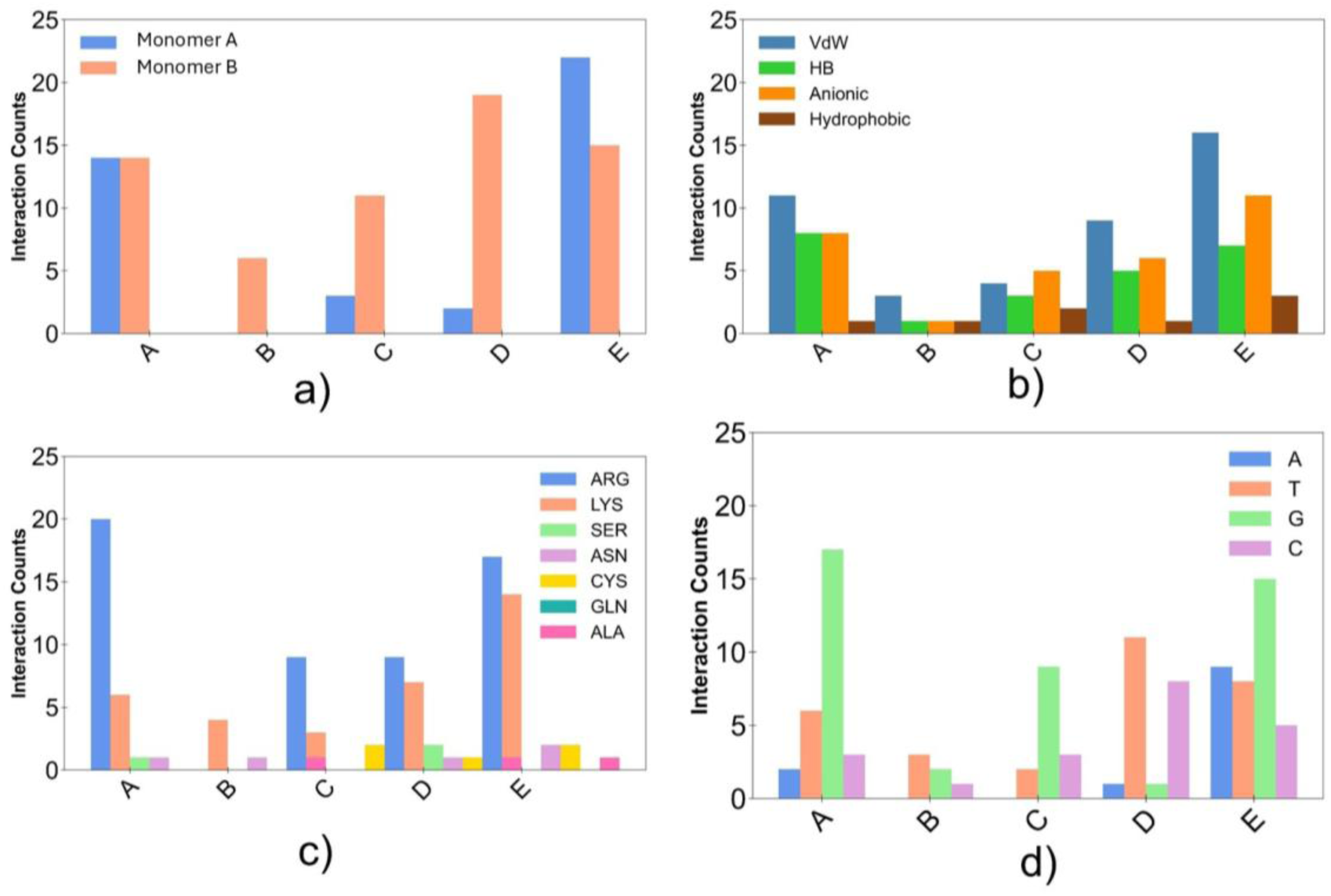
The plot shows the total interaction between ZTA protein and DNA lasting for more than 70 % of the total simulation time of 500 ns. (a) Total interaction counts summing over all kinds of various interactions between monomer A and B of ZTA protein and DNA for all five different systems. (b) Total interaction counts of various interactions like VdW, HB, anionic and hydrophobic interactions for all systems. (c) Total counts of various dominant protein residues participated in the interaction with DNA over the simulation time. d) Total counts of dominant nucleic bases from dsDNA participated in the interaction with ZTA protein for all five systems.

#### a. Various types of interactions

From interaction fingerprint analysis, we computed various interactions, VdW, hydrogen bonds (HB), anionic and hydrophobic contributing to the ZTA-DNA binding (shown in Figure 9(b)). System A shows highest count of VdW inetractions with an equal number of hbond and anionic interactions, and a hydrophobic interaction. In system B, total interactions drop drastically to a very low numbers. System C exhibits moderate recovery of interactions, especially in Hbonds than system B. In system D, interaction count increases signifuicantly, with a notable increment in hbonds and ionic interactions. Among all systems, system E shows highest number of VdW and anionic contacts with lower hbonds than wild type. Although VdW interaction count is higher in comparison to Hbonds, strength of hbonds is higher than VdW interactions. In a recent study by Duraisamy et al. [21], details of interaction strengths for various types of interactions are provided.

#### b. Key ZTA residues

Figure 9(c) shows key amino acid residues from ZTA for various systems. In system A, arginine residues at the DNA binding domain plays crucial role in majority of the interactions. In system B, lysine(K) residues at the DNA binding domain form few interactions with DNA. In system C, two arginine residues from each monomer has been mutated. Remaining arginines at the DNA binding domain forms interaction including a few adidtional interaction by lysine residues maintaing a partial DNA recognization. In system D and E, arginine and lysine residues form major interactions. Hence, arginine at the DNA binding domain is key ZTA residue forming majority of the interaction. It is in good agreement with our previous study[21]. In absence of arginine or in presence of less number of arginine at the DNA binding domain, lysine residues try to counter-balance the interaction load through interaction network rewiring and maintain structural stability of the ZTA-DNA complex.

#### c. Crucial DNA bases

Figure 9(d) shows dominant nucleic bases interacting with ZTA. In system A, guanine (G) is frequently involved in interaction. In system B, guanine was mostly absent and other DNA bases mediate interactions. In system C, guanine and cytosine; in system D,thymine and cytosine form major interactions. In system E, majority of the interactions were mediated by guanine. However, adenine, thymine, and cytosine also formed many interactions. Hence, guanine is crucial nucleic base in the interaction mechanism with ZTA TF.

### Network rewiring of hydrogen bonds

As hydrogen bond is one of the most important interactions in providing stability to the ZTA-DNA complex, we calculated hydrogen bonds contact maps for all systems (Figure 10). System A shows a symmetric hydrogen bonds network involving both monomers in ZTA. The hbonds contact map pattern is like the total-contact map for the wild-type system. Arginine residues from monomer B align along strand I (bases A1-C13) forming majority of the DNA recognizing hydrogen-bonds, particularly residues: B_R179, B_R183, B_R187, and B_R190 form stable hydrogen bonds with respective DNA bases. Among these, B_R183 and B_R187 show continuous occupancy, forming a dense band across multiple neighbouring bases, indicating their key role in stabilizing DNA backbone through direct hydrogen-bonds. In monomer A, the hydrogen-bond network is complementary and concentrated on strand II (bases C14–A26). Residues A_R181, A_R183, A_R187, and A_R190 participate in a series of contacts, forming a pattern that mirrors the B-side arginine across the duplex. A_K178, A_Y180, and A_K194 exhibit broader, moderate-intensity patches, indicating lysine playing an important role in complex stabilization. Their hydrogen bonds are less base-specific but more continuous along the phosphate backbone, helping to maintain electrostatic complementarity and stabilize the overall orientation of the protein on the DNA surface. On the B side, B_K178 shows similar behaviour—wider, less-localized patches, consistent with transient yet recurrent backbone hydrogen bonds. Altogether, the hydrogen-bonds map for system A shows a highly cooperative interaction pattern: arginines provide a strong, localized, and persistent hydrogen bonds responsible for specific recognition on each strand, while lysine reinforces these contacts through broader, stabilizing interactions along the backbone. These combined effect of two residue groups maintain a symmetric and well-anchored ZTA–DNA interface in the wild-type complex.

**Figure 10:**
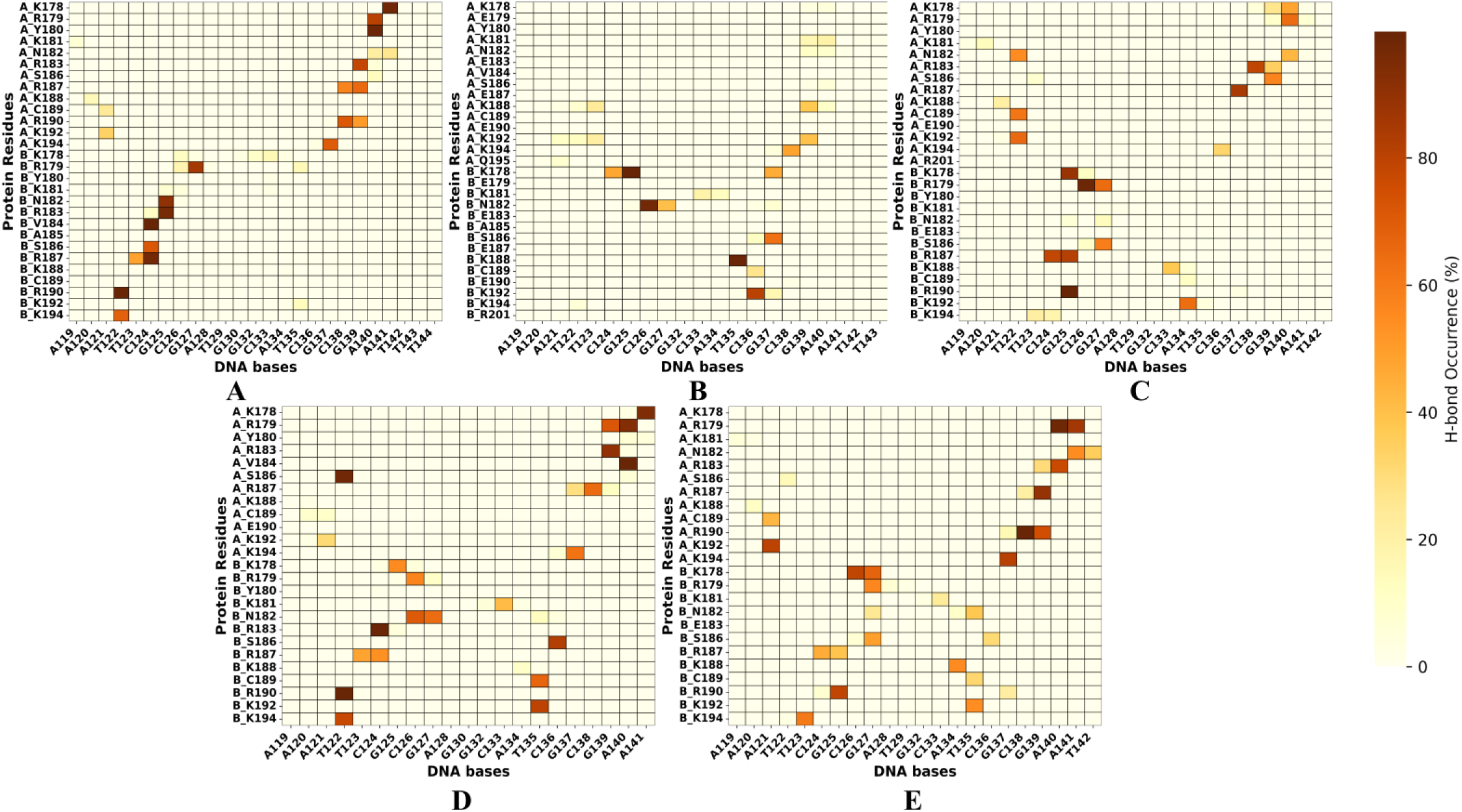
Hydrogen bond frequency maps for five different protein–DNA complexes (A–E), each representing distinct mutations within the DNA-binding domain. The color intensity corresponds to the percentage of hydrogen bond occurrence during molecular dynamics simulations. The wild-type system (A) shows a more continuous hydrogen-bonding network along the major groove, while the mutant systems (B–E) exhibit altered or reduced contact patterns, indicating mutation-induced rearrangements in protein–DNA interaction interfaces.

System B hydrogen-bond contact map shows reduced hydrogen-bond density and continuity in comparison to the wild type. The distinct and well-connected bands of arginine-mediated hydrogen bonds observed in system A are now largely fragmented. In monomer B, where all arginines were replaced by glutamic acids, only a few scattered hydrogen-bond signals remain. The weak and intermittent patches visible near B_K192, B_C189, and B_S186 indicates transient and short-lived hydrogen bonds with DNA backbone. B_K188, B_K192, B_K178, and B_N182 residues form few persistent hydrogen bonds on strand I. Absence of continuous hydrogen-bond band along the core motif region and presence of only isolated signals demonstrate the loss of the extended network, a characteristic feature of the wild type. A similar hydrogen bonds loss was observed in monomer A as well, except few contacts formed by A_K192, A_K194, and A_K188 with low occupancy. The resulting network is asymmetrical and dominated by small, isolated contact points rather than continuous inter-strand hydrogen-bonds.

In system C, a reorganized, yet continuous interaction pattern was observed with several new and redistributed contacts appearing across both strands. In monomer A, residues K178, R179, R183, S186, R187, C189, and K192 form major hydrogen bonds with strand II bases. Among these, R183 and R187 display stronger and extended hydrogen-bonding, while C189 contributes in cross-strand contacts to strand I. Residue K192, which was interacting with strand I in wild-type, now shift its interaction to strand II, showing a consistent contact near the DNA terminal bases. In monomer B, loss of hydrogen-bonding from R183 is accompanied by a compensatory interaction from R179 and R187 on strand I. Importantly, B_R179 emerges as one of the most dominant hydrogen-bonding residues, maintaining a strong and stable interactions across multiple strand I bases. B_K178 also preserves persistent hydrogen bonds near the same region, helping to sustain the backbone connectivity with a few transient interactions by B_K192 and A_C189. Altogether, these changes indicate that while key arginine-mediated hydrogen bonds are diminished at the mutation sites, neighbouring residues, particularly R179, S186 and K192, adopt an extended or shifted roles, producing a rewired yet stable hydrogen-bonding network that maintains ZTA–DNA binding.

In system D, where residue A_R190 is mutated to glutamic acid, the hydrogen-bond network shows a clear redistribution of contacts between two monomers. Unlike the wild type, monomer A now forms hydrogen bonds with both DNA strands, though its overall interaction strength is reduced around the mutation site. The mutated residue A_E190 no longer contributes to hydrogen-bonding, while neighbouring residues—particularly A_S186, A_R187, A_V184, A_R183, A_K178, A_R179— form multiple short hydrogen-bonds across strands. Among these, A_S186 consistently maintain a strong interaction with strand I. In comparison, monomer B carries majority of the overall hydrogen-bonding load and shows extended interaction with strand II, in addition to its usual contacts with strand I. Residues: B_R179, B_S186, B_R190, B_K192, B_K194 form prominent hydrogen-bond patches covering a wide range of DNA bases, while B_C189, B_K178 contribute to additional stable interactions. Overall, system D retains ZTA-DNA complex stability through cooperative redistribution of interaction network: the loss of hydrogen-bonding at A_R190 is compensated by neighbouring A-side residues and by multiple stable contacts from monomer B, which now interacts intensively with both strands, especially strand II.

In system E, where residue B_R183 is substituted, the mutated site (B_R183E) does not form any hydrogen bond with DNA, leaving a clear gap in the interaction pattern on strand I. Nearby residues: B_R179, B_N182, B_S186, B_R187, B_R190 display only weak or intermittent hydrogen bonds with a few short-stable interactions by B_K178 and B_K192. In contrast, monomer A exhibits stronger and extensive hydrogen-bonding, especially with strand II. Prominent hydrogen bonds are observed for A_R179, A_R183, A_R187, and A_R190, forming a continuous column of contacts across several adjacent bases. Among these, A_R190 establishes the most consistent hydrogen bonds, followed by A_R187 and A_R183, all contributing to a partial stable interface along strand II. Additional contacts are formed by A_C189, A_K192 and moderate by A_K178, maintaining a connection near the DNA entry region. Overall, the hydrogen-bond map for system E shows that the loss of contact at B_183 leads to a clear shift of hydrogen-bonding towards monomer A. This redistribution results in a rewired and asymmetric hydrogen-bonding network with monomer A becoming the principal contributor to DNA binding at the absence of the original B_R183 interaction.

The hydrogen-bond analysis elucidates structural and interaction-level observations, revealing how ZTA adapts to mutations while maintaining DNA association. In the wild-type, system A, a dense and symmetric hydrogen-bond network is formed primarily by arginine residues ensuring a balanced contact from both monomers. It produces a compact and stable conformational ensemble. The multiple-mutant system (B) loses this conformational compactness providing a fragmented, short-lived hydrogen bonds and high conformational dispersion, consistent with the observed structural instability and weaker overall binding. The double mutant (C) partially restores hydrogen bonding through compensatory interactions — residues such as A_S186, A_R187, A_C189, and A_K192 form new or extended hydrogen bonds, while B_R179 becomes a dominant contributor. These rewired interactions correspond to the moderate flexibility and maintain a global structure reflected in RMSD and PCA results.

In the single-point mutants, adaptive compensation becomes even clearer. When the mutation occurs in monomer A (system D: A_R190 → E), hydrogen-bonding shifts toward monomer B, which now interacts with both strands and stabilizes the duplex through expanded contacts from B_R179, B_S186, and B_K192. Conversely, when the mutation occurs in monomer B (system E: B_R183 → E), the interaction load transfers to monomer A, whose arginine residues (A_R179, A_R183, A_R187, A_R190) form continuous, long-lived hydrogen bonds across strand II, producing the strongest interaction network among all systems. This cross-monomer compensation mirrors the trends seen in total interaction counts and structural analyses, confirming that ZTA preserves duplex binding through cooperative redistribution rather than rigid dependence on specific residues.

Overall, the hydrogen-bond results demonstrate that mutations at key arginine sites reorganize, rather than eliminating, the interaction network. Lysine and neighbouring residues extend their roles, and the unmutated monomer increases its participation to balance the loss. When correlated with structural properties, like RMSD, RMSF, and PCA data, altogether these findings depict a resilient binding interface with ZTA maintaining its structural stability and DNA recognition through an adaptive hydrogen-bond rewiring and monomeric counter-balancing mechanism.

### Network rewiring: a quantitative perspective based on per-residue binding energy

Further to correlate the described network rewiring and counter-balancing mechanism with residue level binding energy for various orders of mutations, we performed residue-level binding energy decomposition using MM/PBSA. From the analysis we aim to explain how individual residues adjust their energetic contribution which facilitates the network rewiring mechanism effectively to retain complex stability.

Figure 11 shows binding energy per residue specifically for positively charged residues like lysine and arginine in the DNA binding domain. As can be seen from Figure 11, in wild type, major contribution is from arginines in the overall binding of ZTA-DNA complex. Although, ZTA has structural symmetry, however, from the load contributions perspective, monomer B shows slight higher contribution. In addition to arginines, we observe lysine residues (K178, K181, K188, K192, K194) from both the monomers to carry partial load in the interaction mechanism.

**Figure 11:**
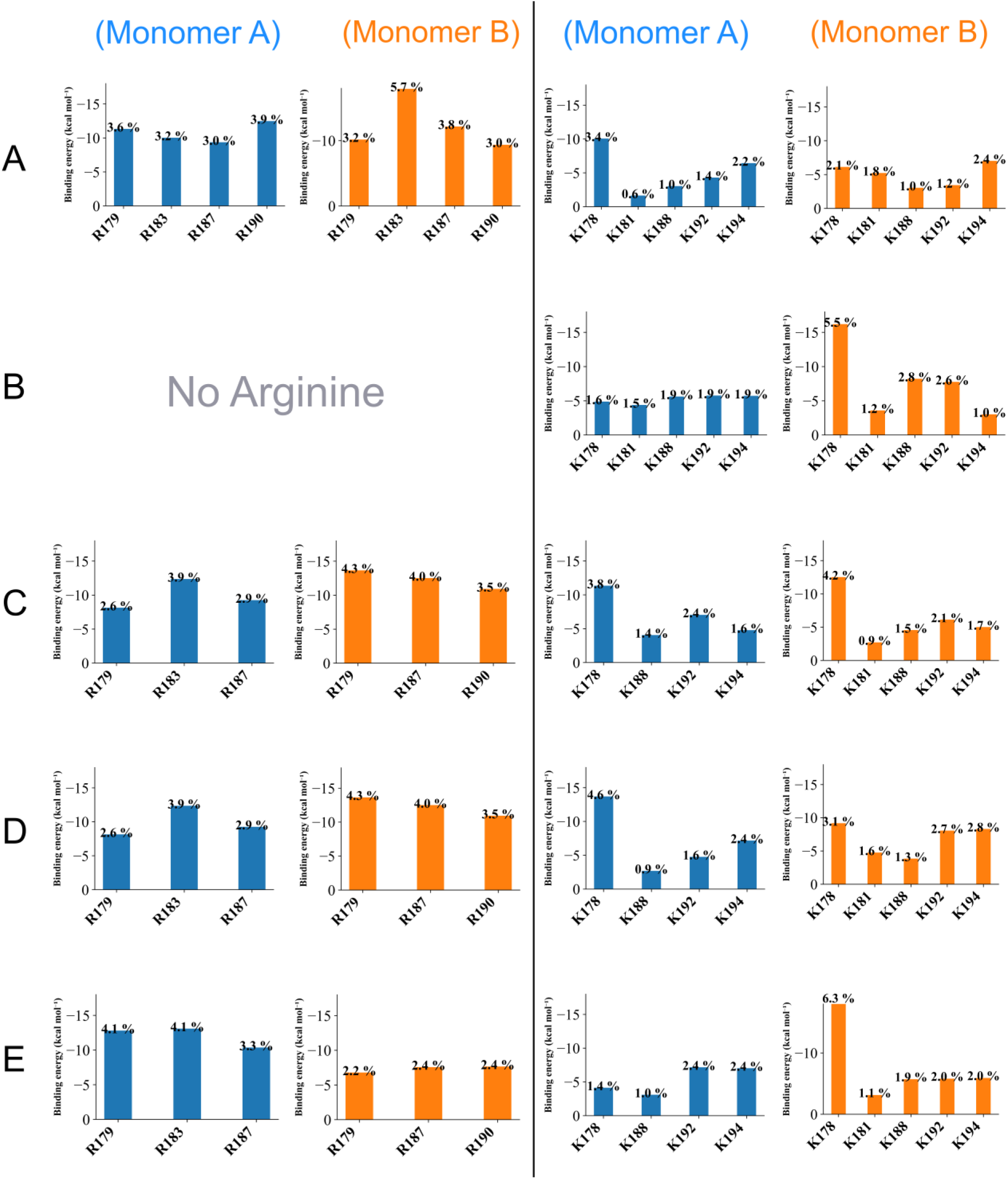
The plots represent the quantitative comparison of load sharing between Arginine and Lysine residues in both A and B monomers of all systems.

In system B, as a result of multi-site mutations, there is a drastic reduction in binding energy. Since arginines at the DNA binding domain have been mutated, nearby lysines are observed to counter-balance the interaction load. From monomer A, lysines contribute 1.5-1.9 % in overall binding. In contrast, monomer B displays a stronger lysine-based compensatory response by K178 (5.5%), K188 (2.8%) and K192 (2.6%) respectively. This dominance of monomer B can be attributed to the fact that monomer B is comparatively closer to the DNA major groove.

System C with double point mutations exhibits a compensatory mechanism, especially monomer B. From monomer B, K178 contributes the most with 5.5% to the binding, followed by K188 and K192, indicating that these lysine residues re-orient to partially share the binding load due to the absence of R183E. In constrast, lysine from monomer A shows minimal contribution to the binding affinity, suggesting that the mutation R190E is not effectively compensated. This assymmetry implies that the structural environment around monomer B allows greater flexibilty and re-organization of lysines to maintain DNA contact, while monomer A remains less accessible.

In system D, with single point mutation in monomer A, lysine and arginine from monomer A contribute around only 21% in the overall binding energy. As A_R190 is mutated, other arginines R179, R183 from monomer A contributes higher percentage of 4% in comparison to wild type. Also K178, K192 show increased contribution than in wild type (system A), especially with K178 (4.6 %). It shows that single point mutation can alter the interaction network in monomer via adjescent residues. However, monomer B does not show much changes except K192 and K194 showing increased contribution in binding.

In system E, with mutation in monomer B at R183E, arginine residues R179, R183, R187, and R190 in monomer A shows higher binding energy contribution with 5.8%, 4.7%, 4.1%, 4.1% respectively, which are maximum among all studied systems. This clearly shows that while R183 is mutated in monomer B, monomer A demonstrate its highest participation in the binding interface. But lysine residues from monomer A shows moderate participation here. Also, residues adjacent to the mutation site showed moderate contribution except K178 from monomer B which showed a contribution of 6.3%, highest among all the systems.

From these quantitative binding energy estimation, we decipher that mutating key sites can alter the interaction network between ZTA and DNA. Especially, in case of single point mutation, we observe certain changes in the interaction mechanism through its opposite monomer association and also with adjacent residues. Lysine residues also plays a crucial role in the binding mechanism particulary in systems C, D and E where only lysines from monomer A participate in the binding.

Overall, the combined energetic and interaction analyses show that ZTA TF-DNA complex try to maintains its structural stability through a compensatory mechanism, dominated by arginine–lysine cooperativity and assymetric cross-monomer re-distribution of the interaction network. This load sharing and counter-balancing mechanism allows the complex to maintain DNA recognition and sustain local perturbations in diverse conditions.

## CONCLUSION

This work demonstrates that the ZTA-DNA complex possesses a highly adaptable interaction framework that allows it to withstand a range of mutational perturbations. Using all-atom MD simulations and binding-energy decomposition, we found that single and double point mutations compensates through a co-operative redistribution of contacts, with nearby arginine and lysine residues from both monomers reinforcing the interface. This compensatory behaviour is especially pronounced in R183E, where the loss of one key interaction triggers broader rearrangements that ultimately increase the total interaction count beyond that of the wild type. Residues such as K178 from monomer B acts as central stabilizers, consistently engaging through hydrogen bonds, electrostatic, and Van der Waals interactions even under substantial stress. In contrast multisite mutations overwhelm this compensatory network, leading to reduced compactness, lower interaction counts, and eventual destabilization of the complex. Previous experimental studies have shown that these mutations can render the protein functionally compromised[13]. Our structural observations provide a plausible explanation for this defect, suggesting that the conformational changes introduced by the mutations may interrupt key steps in the overall process, including the transition toward the lytic state. At the same time, the present analysis highlights how, even in the presence of such defective mutations, the system reorganizes its interaction network to preserve stability. This adaptive rewiring mechanism may represent an important compensatory strategy that allows the system to maintain functional integrity despite the mutational burden.

Overall, our findings indicate that ZTA relies on an adaptive, co-operative network of positively charged residues rather than structural rigidity to maintain DNA binding. This resilience likely represents an evoluionary strategy that enables viral transcription factors to retain function under mutation pressure. Understandng this interaction-rewiring mechanism not only clarifies how ZTA remains operational despite perturbations but also provides insights for designing inhibitors targeting these flexible interfaces and for engineering biomolecules with improvered mutation tolerance. Thus, our rigorous mechanistic studies provide deeper insights on the interaction network reorganization mechanism in ZTA-DNA system which we speculate to be similar for other BZIP family of scissors type of protein TF – DNA systems.

## Data and Software Availability

The initial crystal structure of protein and DNA was taken from the PDB data bank (PDB ID: 2C9N). To develop ZRE3 (Zta responsive element 3) structure we used UCSF Chimera(https://www.cgl.ucsf.edu/chimera/). We used VMD (https://www.ks.uiuc.edu/Research/vmd) and PyMOL (https://www.pymol.org/) for visualization purpose. To perform all-atom molecular dynamics simulation we used GROMACS-2022 software packages (https://manual.gromacs.org/2022/). We performed interaction fingerprints analysis using online available tool ProLIF(https://github.com/chemosim-lab/ProLIF). We performed binding energy decomposition analysis using MMPBSA technique (https://valdes-tresanco-ms.github.io/gmx_MMPBSA/dev/). Initial system coordinates, force field parameters, input scripts for MMPBSA analysis along with generated output, and python scripts for interaction analyses are provided in the following link (https://github.com/dpramanik-lab).

Various other relevant details are provided in the supplementary information.

## SUPPLEMENTARY INFORMATION

Supplementary information provide the following: Details of Zta protein and its missing residues’ modeling; RMSD and RMSF analyses; details of various interaction criterion; interaction fingerprints (Table S1); time series data of hydrogen bonds (Figure S1(a-e)).

## ACKNOWLEDGMENTS

BD and DP thank SRM University – AP for providing HPCC supercomputing facilities. DP thanks Science and Engineering Research Board (SERB), Government of India for supporting with the State University Research Excellence (SURE) project No SUR/2022/004576. BD thanks Anuj Kumar for help in the analysis.

